# Identification of new KCNT1-epilepsy drugs by *in silico,* cell and *Drosophila* modelling

**DOI:** 10.1101/2025.05.22.655336

**Authors:** Michael G. Ricos, Bethan A. Cole, Rashid Hussain, Grigori Y. Rychkov, Zeeshan Shaukat, Nadia Pilati, Stephen P. Muench, Katie J. Simmons, Leanne M. Dibbens, Jonathan D. Lippiat

## Abstract

**Objective:** Hyperactive KCNT1 potassium channels, caused by gain-of-function mutations, are associated with a range of epilepsy disorders. Patients typically experience drug-resistant seizures and in cases with infantile onset, developmental regression can follow. KCNT1-related disorders include epilepsy of infancy with migrating focal seizures and sleep related hypermotor epilepsy. There are currently no effective treatments for KCNT1-epilepsies, but suppressing over-active channels poses a potential strategy.

**Methods:** Using KCNT1 channel structural data, we *in silico* screened a library of known drugs for those predicted to block the channel pore to reduce the current amplitude and inhibit channel activity.

**Results:** Eight known drugs were investigated *in vitro* for their effects on patient-specific mutant KCNT1 channels, with four drugs showing significant reduction of K^+^ current amplitudes. The action of the four drugs was then analyzed *in vivo* and two were found to reduce the seizure phenotype in humanized *Drosophila* KCNT1-epilepsy models.

**Interpretation:** This study identified two known drugs, antrafenine and nelfinavir mesylate, that reduce KCNT1 channel activity and reduce seizure activity in whole animals, suggesting their potential use as new treatments for KCNT1-epilepsy. The sequential *in silico*, *in vitro* and *in vivo* mechanism-based drug selection strategy used here may have broader application for other human disorders where a disease mechanism has been identified.

## INTRODUCTION

Mutations in *KCNT1* are associated with a range of drug-resistant epileptic and developmental neurological disorders. These include severe forms of epilepsy with onset in infancy, including epilepsy of infancy with migrating focal seizures (EIMFS) and later onset focal epilepsies, including sleep related hypermotor epilepsy.^1-4^ The KCNT1 potassium channel subunit is expressed widely in the central nervous system and is activated primarily by intracellular sodium and weakly by depolarization.^5, 6^ *KCNT1* mutations are dominantly-acting heterozygous missense changes in the KCNT1 potassium channel subunit and result in increased channel activity. Accumulating evidence indicates that reduced excitability of inhibitory neurons may be involved in hyperexcitability and seizures.^7-12^

Since the molecular basis of *KCNT1*-associated disorders involves increased KCNT1 channel activity in the central nervous system, its suppression is the basis of stratified therapeutic approaches. Until recently, the only pharmacological agents known to inhibit KCNT1 channels were the non-selective cation channel inhibitors quinidine, bepridil, and clofilium,^13, 14^ each of which have potent effects on the cardiac action potential. Quinidine, which remains in clinical use as a class 1a antiarrhythmic, has been assessed in *KCNT1*-associated disorders, but dosing is limited by its inhibition of cardiac ion channels and dangerous effects on the heartbeat.^15^ Because of this, attempts have been made to identify novel KCNT1 inhibitors that are more potent and selective over other ion channels.^16-18^ Although they are yet to reach clinical use, the efficacy of KCNT1 inhibitors in suppressing electrical activity in mouse models of epilepsy is particularly encouraging.^17-20^ Clinically, there have been case reports of KCNT1 epilepsy patients whose seizures have been reduced with the antitussive drugs tipepidine and dextromethorphan,^21^ the former of which has since been found to inhibit KCNT1 channels^20^, and by fluoxetine, which in addition to several other ion channels has now been found to inhibit KCNT1^22, 23^.

Drug repurposing is the identification of new uses for existing medicines and is particularly attractive for rare genetic disorders as it is associated with significantly lower developmental costs and expedited route to clinical use.^24, 25^ Known drugs that are being assessed for new indications have substantial clinical data, including their safety profile. We therefore hypothesized that existing drugs could be identified that have previously unknown KCNT1 channel inhibition as an off-target effect and that computational techniques could assist their identification. We have previously utilized virtual high throughput screening to identify compounds predicted to occupy the pore in the structure of the chicken KCNT1 channel, generated using cryogenic electron microscopy (cryo-EM).^16, 26^ Since this approach successfully identified novel inhibitors of the human KCNT1 channel,^16^ we aimed to replicate this using a compound library containing known drugs. Here, we describe KCNT1 inhibition as a novel property of four drugs and demonstrate their efficacy with *in vitro* and *in vivo* model systems. In both cellular and *Drosophila* models, we have used several common mutations found in patients with *KCNT1* epilepsy, involving different functional domains of the channel. Thus, these drugs may potentially be effective for a range of patients with different clinical phenotypes.

## MATERIALS AND METHODS

### Virtual screening and drug selection

The intracellular pore region of the chicken KCNT1 channel cryo-EM structure (PDB:5U70)^26^ assigned for docking studies was based upon the predicted binding of other inhibitors.^16^ A 25 Å clip of the structure around these residues was termed as the receptor for docking studies using Glide (Schrödinger Release 2020-2: Glide, Schrödinger, LLC, New York, NY, 2020).^27^ The KCNT1 PDB file was prepared using the Protein Preparation Wizard in the Schrödinger Maestro Graphical User Interface (GUI). This removed steric clashes of amino acid side chains and optimized the position of hydrogen atoms to facilitate docking studies. The academic version of the DrugBank library of known drug molecules (https://www.drugbank.com/academic_research) was downloaded and prepared using the OMEGA module^28^ of OpenEye software (OMEGA version 2.5.1.4 OpenEye Scientific Software, Santa Fe, NM) to produce energy-minimized 3D structures before importing into the Maestro GUI. Glide standard precision screening mode was used to predict the binding pose of each ligand. Drugs were ranked based on predicted binding affinity, likely membrane permeability, and commercial availability.

### Molecular biology

To replicate the heterozygous nature of *KCNT1*-associated disorder and obtain channels comprising wild-type (WT) and mutated subunits, a concatemeric approach was taken. To generate human KCNT1 concatemers, plasmids termed donor and recipient were generated from the pcDNA6-KCNT1 construct used previously^16^ and empty pcDNA6 V5/His6 vector (Invitrogen) using standard polymerase chain reaction (PCR) and cloning techniques. The insert of the donor construct comprised the KCNT1 coding sequence that lacked a start codon and preceded by bases encoding a GGGSGGGS linker. A second donor construct was generated by mutagenesis of this construct, replacing the linker sequence with “self-cleaving” T2A motif from the *Thosea asigna* virus (GSGEGRGSLLTCGDVEENPG),^29^ which exhibits efficient cleavage in CHO cells.^30^ The recipient construct was generated by deleting the stop codon and introducing a unique XhoI site. Sequences containing disease mutation Y796H were subcloned into these constructs from a plasmid used previously.^16^ All sequences generated by PCR were confirmed by Sanger sequencing (Genewiz, Takeley, UK). Finally, the concatemeric construct was generated by subcloning the XhoI/AgeI fragment from the donor plasmid, containing the in-frame linker and complete subunit sequence into the same sites in the recipient plasmid. Recombination-deficient competent *E. coli* cells and 32°C incubation temperature were used for this step to reduce the frequency of deletions between identical sequences in the plasmid, which was confirmed by restriction analysis. For other experiments, constructs with WT, G288S, R398Q or R928C mutant forms of KCNT1 cDNA, tagged with YFP-6His in the pCMV-entry vectors, were created as described previously.^10^

### Cell culture and transfection

Chinese hamster ovary (CHO) cells were cultured in Dulbecco’s modified Eagle’s Medium (ThermoFisher, UK) supplemented with 10% (v/v) fetal bovine serum, 50 U/ml penicillin and 0.05 mg/ml streptomycin, and incubated at 37°C in 5% CO_2_ atmosphere. For whole cell recording of WT/Y796H KCNT1, CHO cells were transiently co-transfected with concatemeric WT/Y796H and EYFP encoding plasmids as described previously.^31^ For electrophysiological experiments, cells were plated onto borosilicate glass cover slips and used 2-4 days later. Human embryonic kidney 293 T cells (HEK 293 T, American Type Culture collection, Rockville, MD) were cultured in the same manner, but with medium supplemented with 2 mM L-glutamine, and 1% (v/v) non-essential amino acids. For excised inside-out recording of homomeric WT or mutant YFP-tagged KCNT1, HEK293 T cells plated on glass cover slips were transfected using Attractene Transfection Reagent (Qiagen, Germany) according to the manufacturer’s instructions and used 2-3 days later.

### Electrophysiology

Whole cell recordings from transiently-transfected CHO cells were conducted at room temperature and analyzed as described previously.^31^ The pipette (intracellular) solution contained, in mM, 100 K-Gluconate, 30 KCl, 10 Na-Gluconate, 29 Glucose, 5 EGTA and 10 HEPES, adjusted to pH 7.3 with KOH and the bath (extracellular) solution contained, in mM, 140 NaCl, 1 CaCl_2_, 5 KCl, 29 Glucose, 10 HEPES and 1 MgCl_2_, adjusted to pH 7.4 with NaOH. For current-voltage and conductance-voltage analysis of the concatemeric constructs, cells were held at -80 mV and 400 ms pulses were applied to voltages between -100 and 80 mV. Conductance values (G) with each voltage pulse (V) were obtained by dividing the peak steady-state current by the driving force (command voltage minus the measured reversal potential). Conductance-voltage data were fit by a Boltzmann function, *G* = (*G_max_* – *G_min_*)/(1+e^(*V*-*V½*)/*k*)^) + *G_min_*, where *G_max_* and *G_min_* are the maximum and minimum conductance values, *V_½_* the half-maximal activation voltage, and slope *k*. The liquid junction potential was calculated and was used to correct voltage values after the data collection. Inhibition by drugs, which were obtained from commercial sources, was determined from currents evoked by 500 ms voltage ramps from -100 to 0 mV at 0.2 Hz as described previously.^16^ Initially drugs (10 mM in DMSO), were diluted to 10 µM in bath solution and applied via gravity perfusion for 2 min, followed by at least 2 min wash with drug-free solution prior to addition of the next drug. Drugs that exhibited current inhibition at 10 µM were analyzed further by concentration-response: *G*/*G_C_* = (1 + ([*B*]/*IC_5_*_0_)*^H^*)^-1^ + *c*, where *G* is the conductance measured as the slope of the current between -60 and 0 mV evoked by the voltage ramp in the presence of the inhibitor, *G_C_* is the control conductance in the absence of inhibitor, [*B*] is the concentration of the inhibitor, *IC_50_* the concentration of inhibitor yielding 50% inhibition, *H* the slope, and *c* the residual conductance.

Inside-out patch clamp recordings were made at room temperature (23^○^C) from transiently-transfected HEK293 T cells as described previously.^11^ The pipette solution contained, in mM, 140 NaCl, 4 KCl, 2 CaCl_2_, 2 MgCl_2_ and 10 HEPES adjusted to pH 7.4 with NaOH. After achieving inside-out configuration, the intracellular face of the membrane was constantly perfused with a solution containing, in mM, 35 NaCl, 110 KCl, 0.2 EGTA and 10 HEPES adjusted to pH 7.3 with KOH. Drug stock solutions of 5-20 mM in DMSO were first diluted in ethanol (1:10) and then in electrophysiological solution. KCNT1 currents were recorded using two voltage protocols, depending on the number of active channels in the patch before drug application. Macroscopic currents (number of active channels >15) were recorded in response to 100 ms voltage ramps between -120 and 120 mV applied every 2 s from a holding potential of -78 mV. The average current amplitudes between 0 and 40 mV were used for the analyses and constructing the concentration-response curves. With a lower number of active channels in the patch (3–15), current traces were recorded in response to 15 s voltage steps to -20, 0 and 20 mV from a holding potential of -78 mV. Currents recorded at each voltage and each drug concentration were averaged over the duration of the trace and normalized to the average amplitudes of the corresponding current traces in drug-free solution. The normalized data for all three voltage steps were then averaged for each drug concentration and used to build the concentration-response curves (see Suppl Fig. 3). To determine the IC_50,_ data were fitted with Hill equation of the form: *Y*=*Bottom* + (*Top*-*Bottom*)/(1+10^((*LogIC_50_*-*X*)\**HillSlope*)), where *Top* was constrained to 1 and *HillSlop*e to -1. Single channel data were analyzed using Ana software developed by Dr Michael Pusch (Istituto di Biofisica, Genova, Italy) (http://users.ge.ibf.cnr.it/pusch/programs-mik.htm).

### Drosophila KCNT1-epilepsy models

The *Drosophila melanogaster* lines carrying the wild type human *KCNT1* transgene (NM_020822_3) or *KCNT1* mutant G288S, R398Q or R928C transgene placed downstream of the yeast UAS promoter, have been described previously.^11^ Wild type or mutant human UAS-*KCNT1* flies were crossed to flies expressing the GAL4 transcriptional activator under the control of the *GAD1* promoter (GABAergic driver, Bloomington stock number 51630 P2) to drive expression of KCNT1 in GABAergic neurons, involved in neuronal inhibition.^11^

### Bang-sensitive behavioral seizure assays in Drosophila

The bang-sensitive behavioral assay (also known as banging assay) was used to test for the presence of a seizure phenotype in *Drosophila*.^11, 32^ Experiments were performed and scored as previously described^11^ with a minimum of 50 *Drosophila* tested for each genotype and drug concentration.

### Analysis of the effects of selected drugs on seizure activity in Drosophila

Based on structural modelling and patch clamping results, four drugs were selected for *in vivo* analysis to determine their effects on the seizure phenotypes in transgenic *Drosophila* lines with human mutant KCNT1 channels. A non-selective inhibitor of KCNT1, bepridil, was included in the analysis for comparison. *Drosophila* food with and without drugs was prepared as previously described.^11^ All chemicals were obtained from Sigma-Aldrich (Gillingham, UK) apart from antrafenine hydrochloride (Toronto Research Chemicals, Canada) and regorafenib (USP, Germany).

*Drosophila* crosses were set up, and the resulting embryos were collected, as previously described (Hussain et al 2024). A range of concentrations of each drug dissolved in *Drosophila* food, between 0.001 µM and 1 µM, were used in the experiments. Multiple replicates were performed for each dosage of the drugs and controls. Controls were Vehicle Control (VC) which was the normal food with just the solvent ethanol present (1 µl/1 ml) in the *Drosophila* food, and Normal Food control (NF) where no drug or solvent (ethanol) was added in the *Drosophila* food.

### Statistical analysis

In all experiments, experimenters were not blinded to variants or treatments, nor were samples randomly assigned to groups. Sample sizes were not calculated in advance and no data were excluded from the analysis. Data were analyzed using GraphPad Prism 9 software (San Diego, CA, USA). All values are reported as mean ± standard error (SE). All data passed normality and lognormality Shapiro-Wilk tests, determined by GraphPad Prism 9. In patch clamp data ‘n’ represents number of cells, and all experiments were repeated using cells from a minimum of two separate transfections. In the experiments analyzing the effects of drugs in *Drosophila*, statistical significance of differences between groups was determined using Brown-Forsythe and Welch’s ANOVA tests (Supplementary Table 3), assuming non-equal SDs, followed by Dunnett’s multiple comparisons tests (GraphPad Prism 9). The results of the Dunnett’s multiple comparisons tests are shown in Supplementary Table 4, where ‘*N*’ represents independent trials using independent fly crosses, with the total number of flies for each condition shown in brackets.

## RESULTS

### Identification and selection of drugs for functional evaluation

Having previously validated the use of chicken KCNT1 structural data in virtual screening to identify novel inhibitors of human KCNT1,^16^ we turned our attention to existing drugs. The predicted binding modes of the Drugbank compound library in the KCNT1 intracellular pore vestibule structure were computed and ranked by their docking score. These were further filtered by their clinical status as known drugs, and manually selected based on their binding mode, commercial availability and computed hydrophobicity (clogP>3). Nine drugs were selected for functional assessment: indinavir, antrafenine, candesartan cilexetil, nelfinavir mesylate, regorafenib, dihydrotachysterol, atorvastatin, lifitegrast, and terconazole.

### Generation of a cellular model of heterozygous assembly of wild-type and Y796H KCNT1 subunits

Several KCNT1 channel inhibitors differ in their potency between channels comprised of wild-type (normal, WT) subunits alone and those formed by subunits carrying a disease-associated gain-of-function mutation.^16, 17, 33^ Given the heterozygous dominant nature of KCNT1-disorders, we aimed to determine KCNT1-inhibiting properties of drugs using channels containing both WT and mutant subunits. Initially, we generated KCNT1 concatemers by fusing two WT subunits with a flexible linker and expressed the construct in CHO cells. This yielded outwardly-rectifying currents that resembled WT KCNT1 (Fig. 1A), but with conductance-voltage relationships shifted to more negative potentials (Fig. 1C). In rodents, different *Kcnt1* mRNA transcripts are generated from alternative transcription initiation sites, leading to variation in the amino terminus of the channel subunit and channels with altered activation kinetics.^34^ We therefore reasoned that constraining the amino terminus of the second subunit in the concatemer may underlie this altered activity and the linker sequence was replaced by a T2A “self-cleaving” motif. Expression of this construct in CHO cells resulted in similar currents, but with activation kinetics more closely resembling WT channels (Fig. 1). The second subunit in this concatemer was then replaced with one harboring the Y796H mutation in the distal C-terminus and used for functional evaluation of drugs predicted to occupy the channel pore. Channels produced from this construct had activation half-maximal voltages intermediate of homomeric WT and Y796H KCNT1 channels. A summary of the activation kinetics of the monomeric and concatemeric channel currents is provided in Supplementary Table 1.

**Figure 1.**
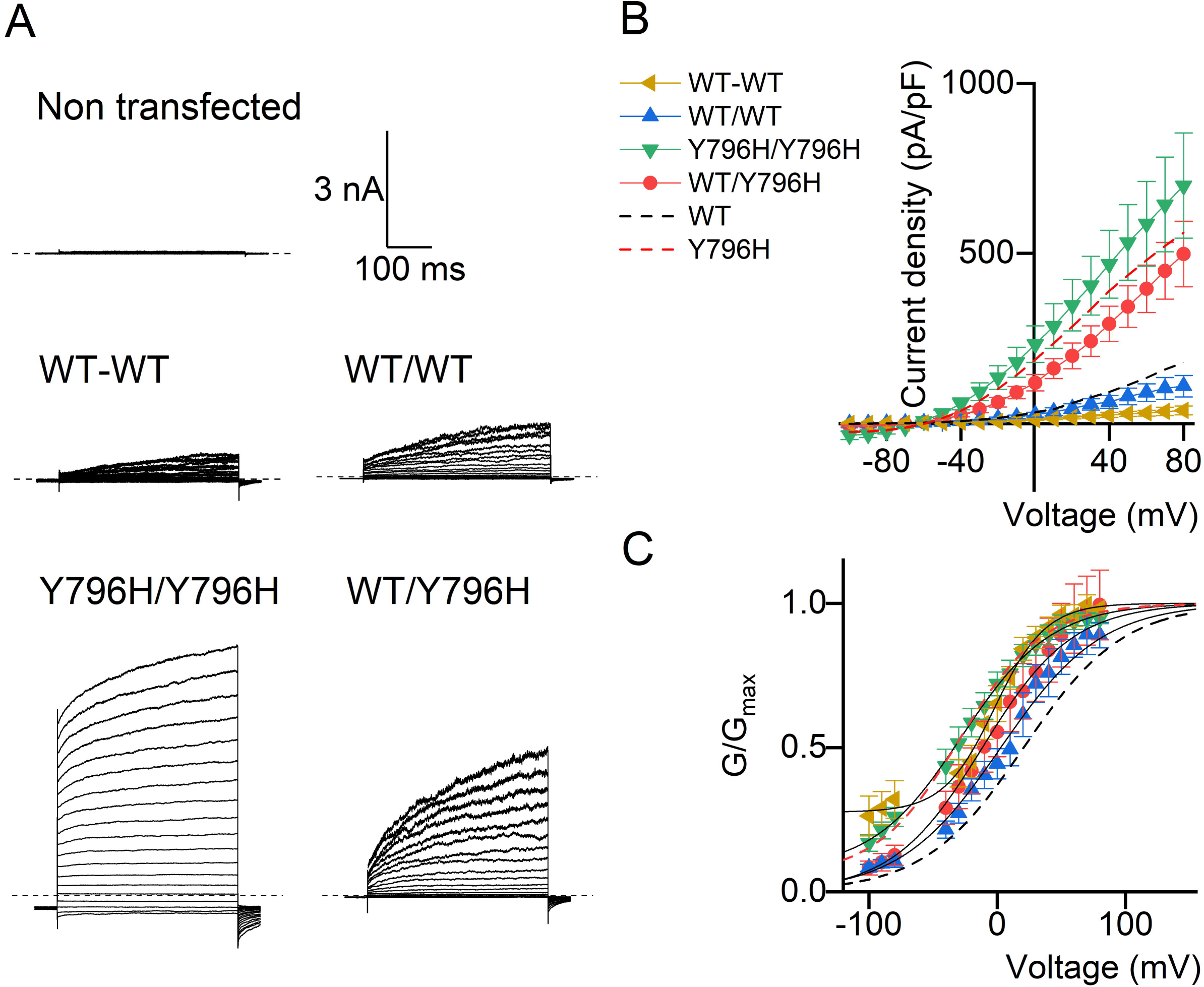
Generation of an expression construct for heteromeric expression of WT and Y796H KCNT1 subunits. The concatemeric constructs encode two KCNT1 subunits, separated by either a tethered EGGGSGGGS motif or a T2A self-cleaving peptide between the first and second subunit. **A** Representative current traces recorded from non-transfected CHO cells using whole-cell patch clamp or with CHO cells transfected with constructs containing WT KCNT1 subunits in both the first and second position (WT-WT with a hyphen indicates the tethered concatemer; WT/WT with a slash indicates the cleavable T2A construct), Y796H KCNT1 in both the first and second positions (Y796H/Y796H), or WT in the first and Y796H KCNT1 in the second positions (WT/Y796H), as indicated. The dashed line indicates the zero-current levels. **B** Current-voltage and **C** Conductance-voltage relationships from the currents recorded. Mean data for monomeric WT KCNT1 (black) and Y796H (red) are indicated as dashed lines for comparison. Data are mean ± SEM (*n*= 6 cells for WT-WT and WT/Y796H; *n*=5 cells for WT/WT and Y796H/Y796H).

### Functional evaluation of drugs in inhibiting WT/Y796H KCNT1 channels

Using the WT/Y796H KCNT1 construct as a model for “heterozygous” *KCNT1* pathogenic variants, the drugs selected from the molecular docking were applied to transfected cells in patch clamp experiments. Four of the drugs, antrafenine, atorvastatin, nelfinavir mesylate and regorafenib inhibited WT/Y796H KCNT1 channels expressed in CHO cells when tested at 10 µM concentration. Concentration-inhibition analysis yielded mean ± SEM *IC_50_* of 1.30 ± 0.2 µM (n=5) for antrafenine, 2.86 ± 0.30 µM (*n*=5) for nelfinavir mesylate, 7.54 ± 0.99 µM (n=6) for atorvastatin, and 10.30 ± 1.25 µM (*n*=6) for regorafenib (Fig. 2A, B). Candesartan cilexetil appeared less potent, and higher concentrations were not attempted as it appeared to require a range at least an order of magnitude higher (Figure 2B). On the other hand, indinavir, dihydrotachysterol, liftegrast, and terconazole did not exhibit inhibition when tested at 10 µM concentration (Fig. 2C) and were not studied any further.

**Figure 2.**
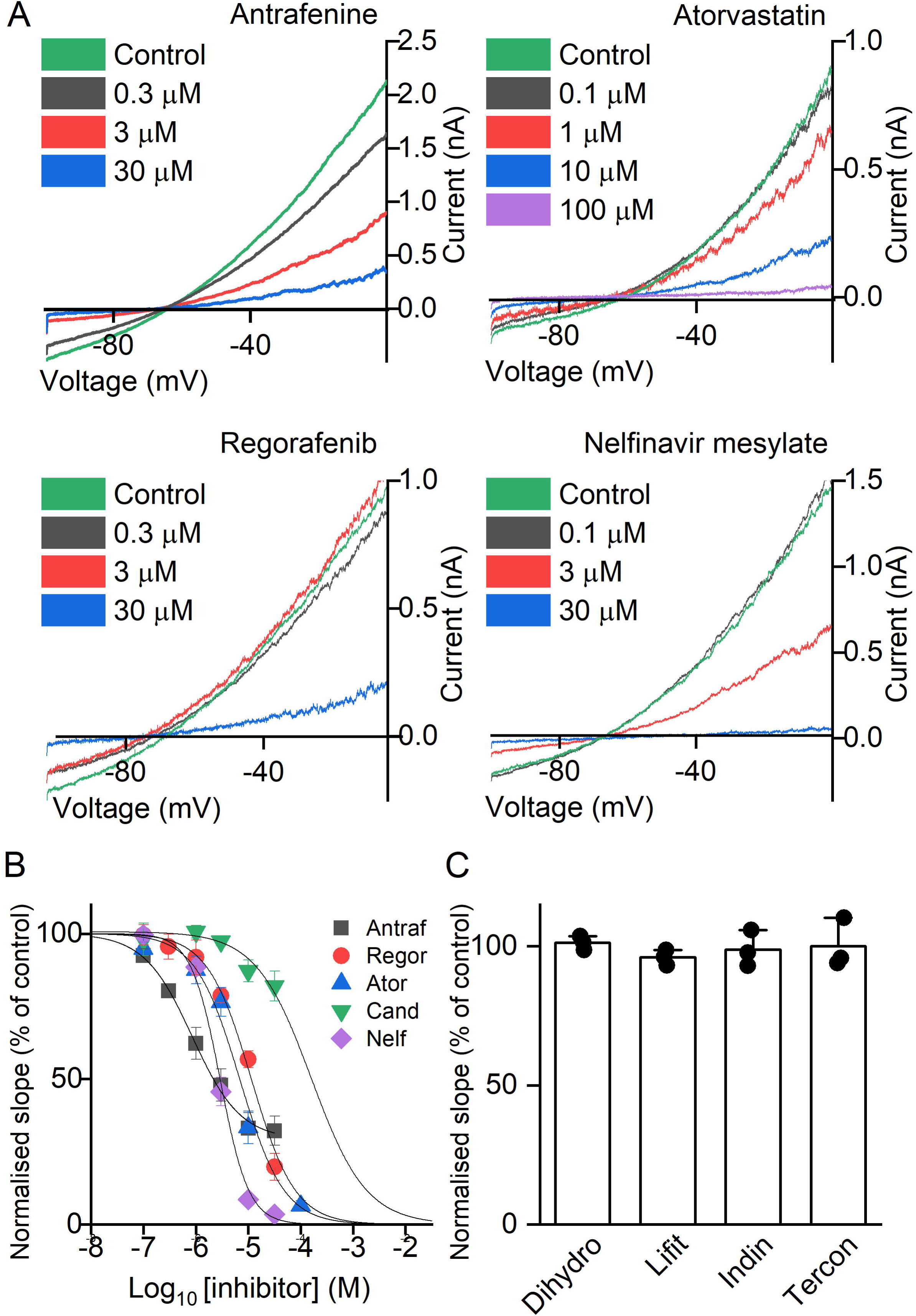
Functional evaluation of known drugs with WT/Y796H KCNT1 channels expressed in CHO cells. **A** Representative traces and **B** mean ± SEM. concentration-dependent inhibition by active inhibitors. Atraf, antrafenine (*n=*5 cells); Regor, regorafenib (*n=*6 cells); Ator, atorvastatin (*n=*6 cells); Cand, candesartan (*n=*5 cells); Nelf, nelfinavir (*n=*5 cells). **C** Drugs inactive at 10 µM. Data are mean ± SEM WT/Y796H KCNT1 conductance measured as the slope of the current evoked by a voltage ramp to from -100 to 0 mV in the presence of 10 µM inhibitor, relative to control conductance prior to drug application. Dihydro, dihydrotachysterol (*n=*3 cells); Lift, liftegrast (*n=*3 cells); Indin, indinavir (*n=*3 cells); Tercon, terconazole (*n=*3 cells).

Similar to the inhibitors that we previously identified by virtual screening,^16^ these drugs were predicted to bind to the intracellular vestibule of the KCNT1 channel pore below the selectivity filter (Fig. 3, Supplementary Fig. 1). Each are predicted to make hydrophobic and hydrogen bond interactions with pore-lining amino acids, here provided as the equivalent amino acid and number in the human KCNT1 homolog. Antrafenine and atorvastatin are both predicted to make a hydrogen bonding interaction with the side chain of T314, which forms the intracellular-facing part of the selectivity filter, with atorvastatin making two additional hydrogen bonding interactions with the E347 side chain in the S6 segment. Nelfinavir and regorafenib are both predicted to hydrogen bond with the backbone carbonyl of F312, with nelfinavir also interacting with the side chain of T314. Antrafenine contains two and regorafenib contains one terminal trifluoromethyl group. Similar to the predicted binding modes of BC12 and BC14,^16^ we found that these terminal groups occupy a cavity formed between adjacent S6 segments and the pore helix. Antrafenine spans the pore, with the two trifluoromethyl groups at the opposite ends of the molecule occupying the equivalent cavities formed by opposite subunits in the KCNT1 tetramer. Since carrying out this docking, structures of human KCNT1 in several states, including with an inhibitor bound were published.^35^ With the exception of atorvastatin, there is good agreement in the docking scores of these drugs between the chicken and inhibitor-bound human KCNT1 structures (Supplementary Table 2), with close conservation of both sequence and structure in the pore domain (Supplementary Fig. 2).

**Figure 3.**
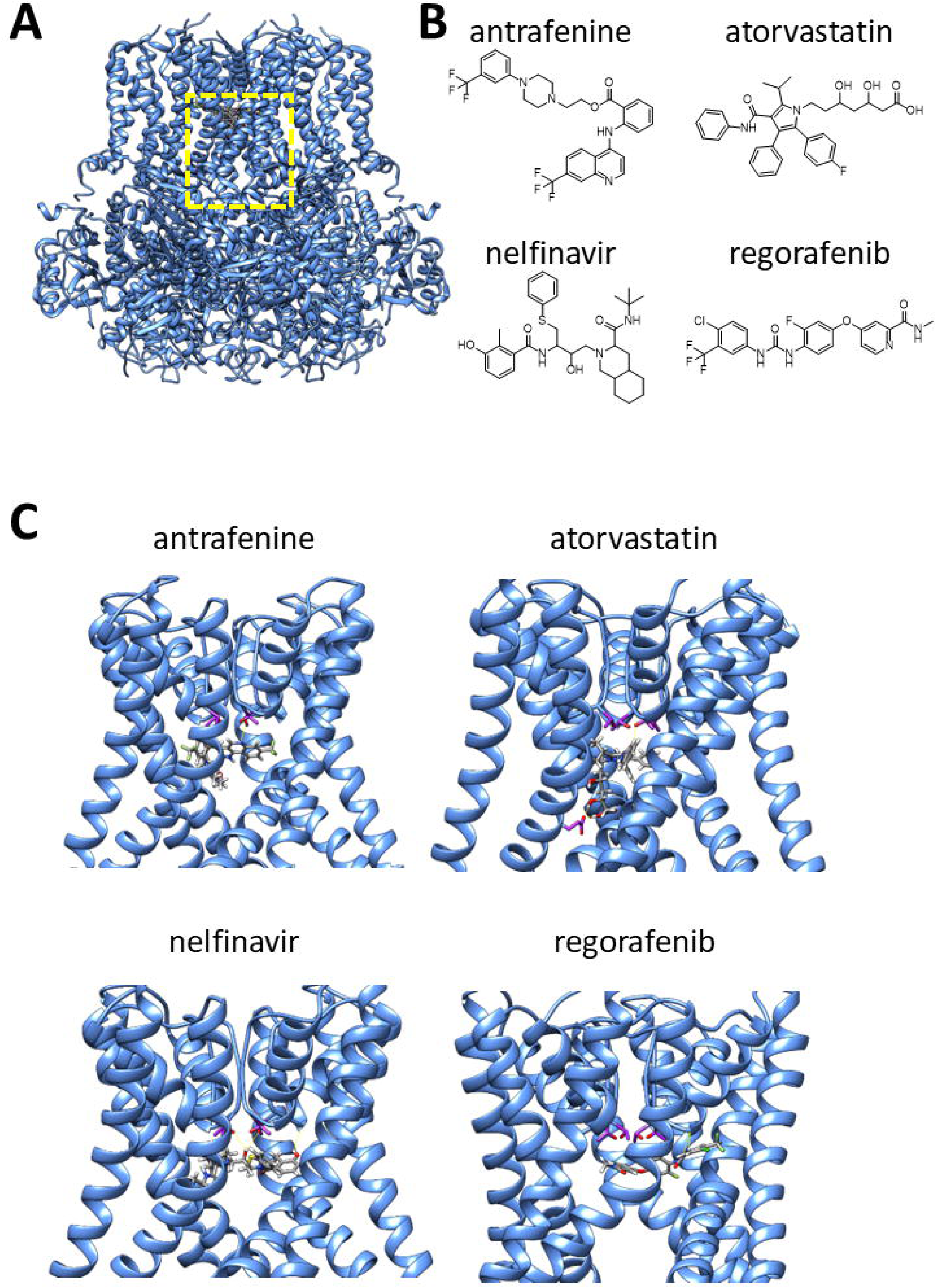
Molecular docking of drugs to the KCNT1 channel pore. **A** Structure of chicken KCNT1 in the active conformation (PDB:5U70)^26^ with drugs docked to the channel pore domain, indicated by the yellow dashed box. **B** Molecular structures of active drugs identified in this study. **C** Individual active drugs docked to the KCNT1 pore domain as indicated. For clarity, only the S5, pore helix, selectivity filter, and S6 of each subunit is shown and rotated to best illustrate each drug binding mode. Threonine sidechains at the intracellular end of the selectivity filter are colored magenta, also the aspartate side chain in the S6 segment that interacts with atorvastatin.

### Inhibition of single human KCNT1 channels by the four prioritized drugs

The four drugs antrafenine, nelfinavir mesylate, atorvastatin and regorafenib were further analyzed by investigating their effects on unitary WT and R928C, G288S and R398Q mutant KCNT1 channels expressed in HEK293 T cells using the inside-out configuration of the patch clamp technique. The tight seal between the patch pipette and the cell was achieved in the control bath solution, after which the inside-out patch was excised and moved under the outlet of the gravity-fed perfusion system used to apply drugs to its intracellular face. Consistent with the prediction that they directly block the pore, antrafenine and nelfinavir were seen to inhibit over 60% of the activity of the WT human KCNT1 channel at concentrations of 50 nM (Fig. 4A, B). The effects of atorvastatin and regorafenib on KCNT1 channels were less potent, with the drugs not fully inhibiting KCNT1 activity at concentrations of 10 µM (Fig. 4C, D). All inhibitors reduced the open probability of KCNT1 without affecting the single channel conductance (Fig. 4, insets). Antrafenine, nelfinavir mesylate and atorvastatin had similar inhibitory effects on WT and each of the R928C, G288S and R398Q mutant KCNT1 channels at the same concentrations (Fig. 5, Table 1, Supplementary Fig. 3).

**Figure 4.**
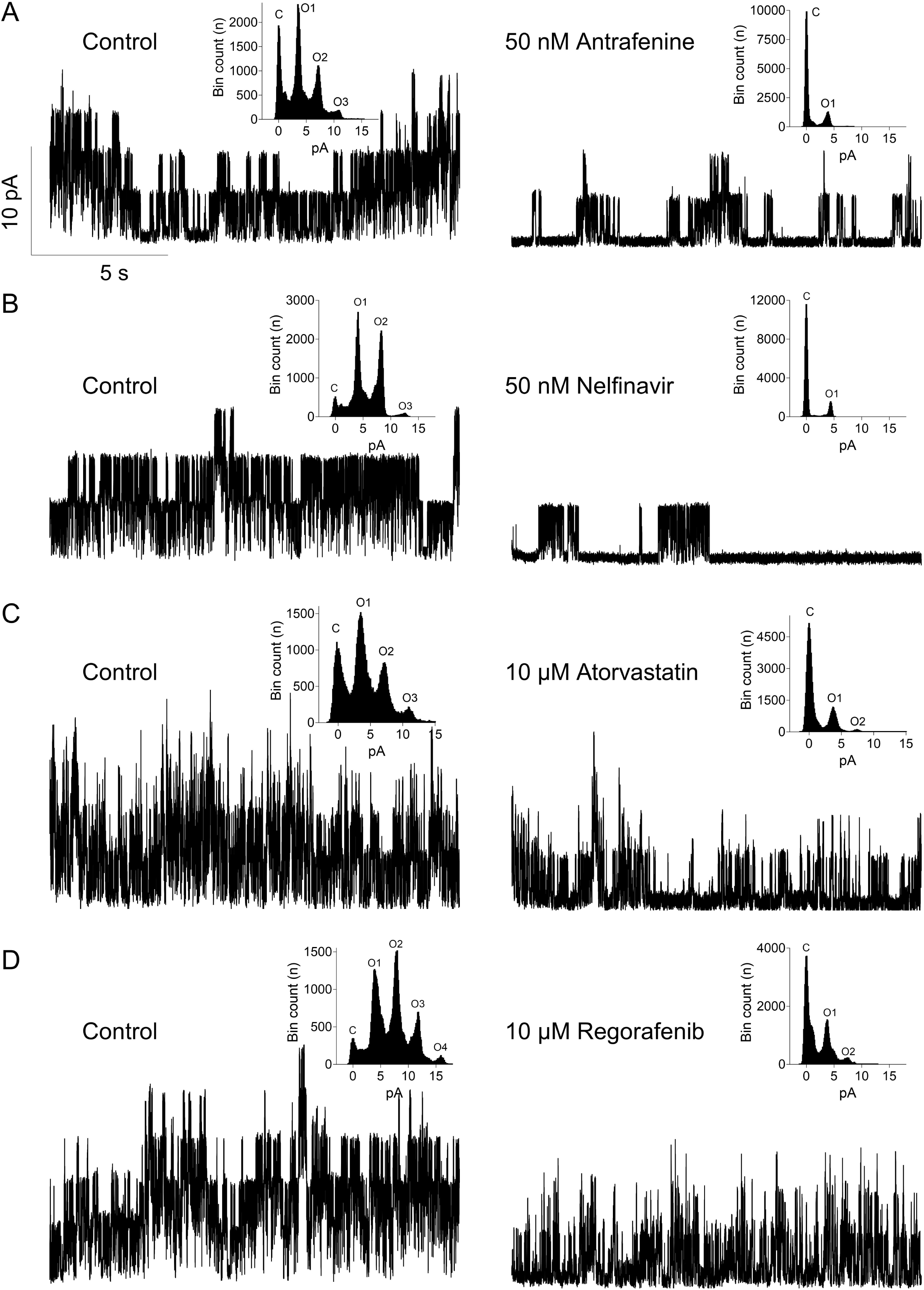
Functional assessment of drug efficacy using inside-out patch clamping of HEK293 T cells expressing WT KCNT1 channels. Single channel current traces were recorded at 0 mV and physiological K^+^ gradient under control conditions and in the presence of different drugs: **A** Antrafenine (50 nM), **B** Nelfinavir mesylate (50 nM), **C** Atorvastatin (10 µM), and **D**. Regorafenib (10 µM). Insets show all-point amplitude histograms of the corresponding traces (C closed and O open levels).

**Figure 5.**
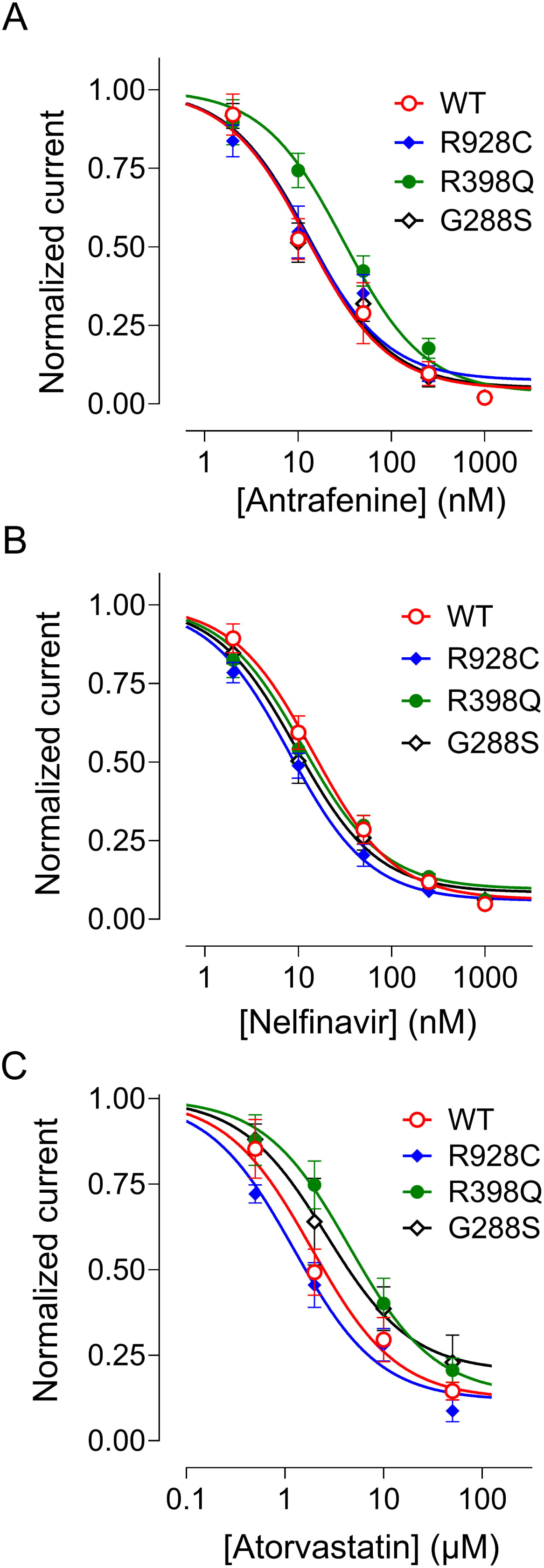
Assessment of known drug efficacy in inhibiting KCNT1 channels carrying epilepsy-causing mutations, G288S, R398Q, and R928C. Concentration-response curves were constructed using KCNT1 currents (see Supplementary Fig. 3) recorded in inside-out patches in the presence of different concentrations of a drug in the bath and normalized to the current amplitude recorded before drug application. The data points were fitted with a Hill equation with a slope of -1. **A** Antrafenine, **B** Nelfinavir, and **C** Atorvastatin. (For *IC_50_* values see Table 1)

**Table 1.** Potency of drugs inhibiting wild-type (WT) and mutant KCNT1 channels in inside-out patches. *IC_50_* values for antrafenine, nelfinavir and atorvastatin-mediated inhibition of WT and mutant KCNT1 channels in inside-out patches, derived from concentration-response curves shown in Fig. 5. Data are mean ± SEM of *<u>n</u>*patches.

| Variant | Antrafenine | Nelfinavir | Atorvastatin |
| --- | --- | --- | --- |
| WT | 12.6 ± 3.6 nM (n=6) | 14.3 ± 2.6 nM (n=6) | 1.9 ± 0.6 µM (n=5) |
| R928C | 12.3 ± 3.9 nM (n=6) | 8.0 ± 1.1 nM (n=5) | 1.2 ± 0.3 µM (n=4) |
| R398Q | 30.7 ± 7.4 nM (n=5) | 11.2 ± 2.2 nM (n=6) | 4.5 ± 1.7 µM (n=5) |
| G288S | 13.3 ± 3.7 nM (n=5) | 9.5 ± 1.8 nM (n=6) | 2.6 ± 1.6 µM (n=3) |

### In vivo analysis of the four prioritized drugs fed to Drosophila models of KCNT1-epilepsy

To analyze the effects of the four prioritized drugs on *KCNT1*-related seizures in an animal model, we used our *Drosophila* models of *KCNT1* epilepsy. The three models contain human *KCNT1* with patient-specific mutations G288S, R398Q or R928C. A *Drosophila* line with WT human *KCNT1* was used as a control. The expression of *KCNT1* was under the control of the yeast UAS promoter and crossing to a line with the GAL4 transcription factor under the control of a chosen promoter enables tissue-specific expression of the transgene.^36^ We have shown previously that expression of mutant KCNT1 channels in GABAergic inhibitory neurons gives a seizure phenotype in bang-sensitive assays and that the animal models can be used to analyze the effects of drugs on KCNT1 channel activity.^11^ Here we used drug-feeding assays to test the effect of the four drugs on the seizure phenotype of the three different *Drosophila* lines carrying a patient *KCNT1*-epilepsy mutation.

Juvenile *Drosophila* (larvae) from the lines expressing WT, G288S, R398Q or R928C human KCNT1 channels, were fed each of the drugs and adult *Drosophila* were analyzed for seizure activity using bang-sensitive assays. Each mutant line was also raised on normal *Drosophila* food (NF) or normal *Drosophila* food containing the vehicle ethanol (VC) which was used to introduce each of the drugs to the *Drosophila* food. These two controls showed the baseline seizure activity of each of the mutant KCNT1 *Drosophila* lines (Fig. 6). As shown in Fig. 6, two of the drugs, antrafenine and nelfinavir mesylate, were seen to significantly reduce the seizure activity in each of the three *Drosophila* mutant *KCNT1* lines in a dose dependent manner (see Supplementary Tables 3 and 4 for statistics). The other two drugs, atorvastatin and regorafenib, did not show a significant effect on seizure activity in the G288S, R398Q or R928C mutant lines. Analysis of 1 µM atorvastatin was excluded due to reduced viability of adult flies at this concentration. Nelfinavir mesylate showed the greatest reduction in seizure activity, followed by antrafenine. The known non-specific inhibitor bepridil was included for comparison and seen to exacerbate the seizure phenotype in each of the three *KCNT1* mutant lines (Fig. 6).

**Figure 6.**
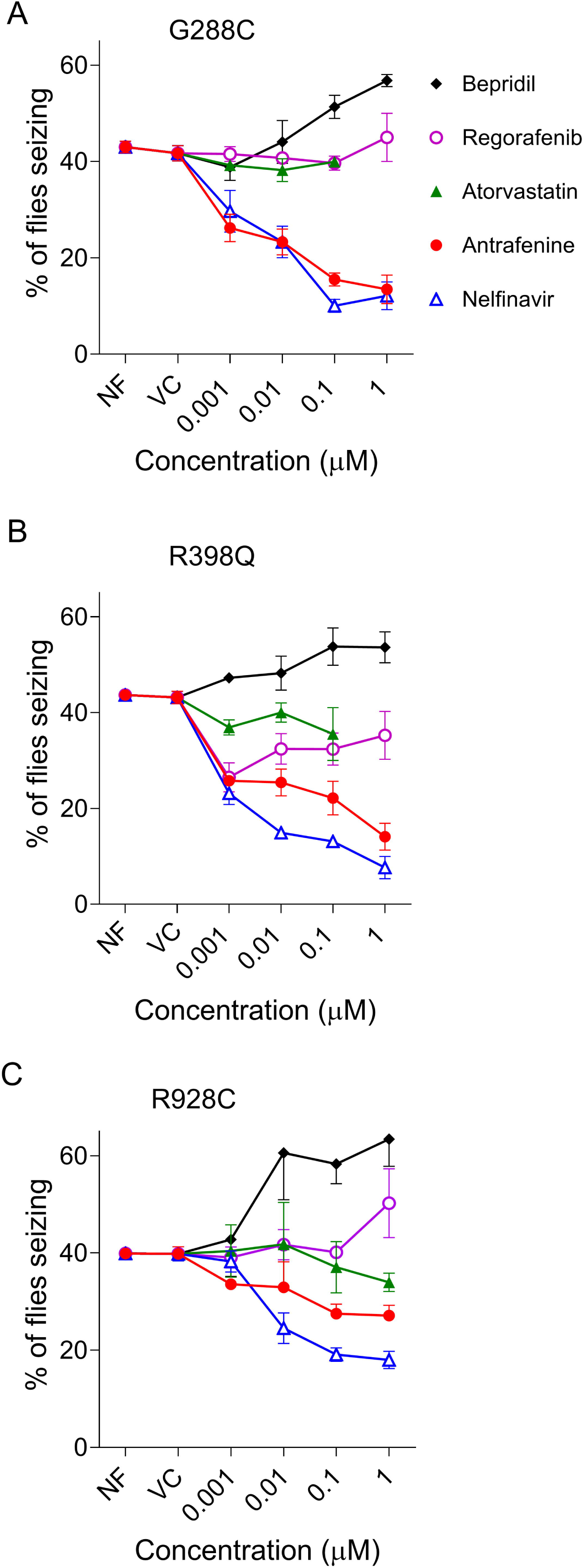
Reduction of seizure phenotype in three *Drosophila* KCNT1 mutant lines by known drugs. The percentage of adult *Drosophila* showing seizure activity is shown for each of the KCNT1 mutant lines G288S, R398Q or R928C when raised on Normal Food (NF) or food with added Vehicle Control (VC), or the known drugs nelfinavir mesylate, antrafenine, atorvastatin or regorafenib. Seizures in *Drosophila* are caused by expression of human *KCNT1* transgenes with mutations in the pore-adjacent loop domain, G288S (**A**), RCK1 domain, R398Q (**B**) or the RCK2 domain, R928C (**C**) in inhibitory GABAergic neurons. Data are presented as mean ± SEM.

## DISCUSSION

We have successfully used *in silico* screening to identify four existing drugs that inhibit KCNT1 channels at low micromolar potency in whole-cell patch clamp recordings, and importantly more potently than is reported for the known ion channel blocker quinidine. The outcomes from this approach are similar to those we obtained previously with a library of commercially available small molecules.^16^ Since the four drugs have already been considered for therapeutic use, pharmacological and clinical data are available to inform whether they can be trailed for treating *KCNT1*-associated disorders. Using inside-out patch clamp recordings, antrafenine and nelfinavir were seen to inhibit 80 - 90% of the activity of the wild type and mutant human KCNT1 channels at 250 nM. This enhanced sensitivity, compared to whole-cell patch clamp, supports the idea that the drugs block the channel at the intracellular pore vestibule of the channel, either via the cytoplasm or directly from the lipid bilayer.

Based on the *in vitro* analysis and their inhibition of KCNT1 channels, the four drugs antrafenine, atorvastatin, nelfinavir mesylate, and regorafenib, were selected to be analyzed *in vivo* for their effects on the seizure phenotype in three *Drosophila* models of *KCNT1*-epilepsy. *Drosophila* was chosen as an animal model as the key components in the regulation of neuronal excitability in humans and *Drosophila* are highly conserved.^37^ Other *Drosophila* models of human genetic epilepsies include those for sodium channel *SCN1A*-related epilepsy^38^ and Pyridox(am)ine 5′-phosphate oxidase (*PNPO*)-related epilepsy.^39^ The relative ease of genetic manipulation, fast generation times and the low cost of housing make *Drosophila* attractive for modelling human epilepsy and for relatively rapid drug analyses. The four candidate drugs were fed to each of our three Drosophila models of KCNT1-epilepsy that contain three of the most common patient missense mutations, G288S, R398Q or R928C. Antrafenine and nelfinavir mesylate were found to significantly reduce the seizure phenotype in each *Drosophila* mutant *KCNT1* line, while atorvastatin and regorafenib did not. Nelfinavir was seen to reduce the seizure phenotype by 50% in *Drosophila* with G288S and R398Q and R928C. Antrafenine reduced the phenotype by at least 50% in G288S and R398Q, with a 25% reduction in R928C. The reduction of the seizure phenotype was seen to be dose-dependent for each of these drugs, showing suppression at 0.001 µM, with the greatest effects at 0.1 µM or 1 µM in food.

The two potential new drugs for treating people with *KCNT1*-epilepsy identified in this study are yet to be trialed in patients for this indication. However, it is possible that our study demonstrates the benefit of employing whole animal models when assessing potential drugs for their effectiveness in reducing seizures in human epilepsy. Non-selective cation channel blocking drugs, such as quinidine and bepridil have been shown to inhibit KCNT1 channel activity in *in vitro* experiments,^33, 40^ leading to quinidine being trialed as a stratified treatment for *KCNT1* epilepsy.^41-44^ However, quinidine has had mixed results in treating patients with worsening of seizures in some patients.^1, 45^ The use of bepridil has also been limited in clinical settings over safety concerns and was withdrawn for this reason.^46^ We have shown that both quinidine^11^ and bepridil (this study) exacerbate the seizure phenotype in the *Drosophila* models of *KCNT1*-epilepsy, most likely by inhibiting other cation channels that control neuronal excitability, which likely highlights the value of *in vivo* pre-clinical analysis for drug re-purposing. Interestingly, seizures were most exacerbated by bepridil in the R928C mutant line, suggesting mutation-specific differences in response to drugs and indicating a role for mutation-specific pharmacogenomics.

Nelfinavir mesylate and antrafenine showed significant reduction in seizure activity associated with all three *KCNT1* patient mutations R928C, R398Q and G288S when tested in our animal models. They also showed significant reduction of K^+^ currents in HEK cells expressing the same three mutant channels, as well as channels comprised of WT and Y796H mutant KCNT1 subunits in CHO cells. These data suggests that these two drugs have the potential to inhibit a spectrum of mutant *KCNT1* channels found in patients and, if found to be clinically effective, could be new treatments for patients with a range of different *KCNT1* mutations. Apart from antrafenine, each of the KCNT1-inhibiting drugs described here are presently in clinical use. Nelfinavir mesylate (brand name Viracept) is used to treat HIV infection, inhibiting the HIV-1 protease. It has been reported that this drug inhibits hERG potassium channels with an IC_50_ of 11.5 µM,^47^ which could place patients at risk of arrhythmia. Atorvastatin (brand name Lipitor) is a competitive inhibitor of 3-hydroxy-3-methyl-glutaryl-coenzyme A (HMG-CoA) reductase, an enzyme involved in cholesterol synthesis, and is a statin used in the treatment of hyperlipidemia and hypercholesterolemia. Partial inhibition of hERG currents by atorvastatin at low micromolar concentrations have been reported and appear to affect hERG channel inactivation kinetics.^48^ Regorafenib (brand name Stivarga) is an anticancer drug that targets receptor tyrosine kinases and is presently under clinical evaluation for treating brain tumors.^49^ Antrafenine (brand name Stakane) has analgesic and anti-inflammatory effects, but there is no record of its clinical availability or use since the mid-1980s. In a clinical evaluation with osteoarthritis patients, mean circulating plasma levels of antrafenine were measured at around 0.3 µM,^50^ which is in the concentration range of KCNT1 channel inhibition reported here. Further evaluation of the pharmacodynamics and pharmacokinetic properties of each of the drugs identified here will be required to justify their clinical evaluation in *KCNT1* epilepsy.

Two out of the four candidate drugs tested suppressed seizures in our whole animal model of *KCNT1* epilepsy, indicating that our combination of *in-silico* screening, *in-vitro* assay and *in-vivo* drug testing approach may be an effective strategy in identifying drugs for repurposing to treat *KCNT1* epilepsy. This combined approach may be applicable to other human disorders where a disease mechanism has been identified.

## ACKNOWLEDGMENTS

Supported by a BBSRC-CASE studentship in conjunction with Autifony Therapeutics Ltd awarded to B.A.C. (BB/M011151/1). We acknowledge support from the Channel 7 Children’s Research Foundation Grant 13435028 (LMD, GYR, ZS), and National and Health Medical Research Council of Australia (Senior Research Fellowship 1104718 and Project Grant 1125523 to LMD, and Ideas Grant 2028987 to LMD, GYR and RH). We thank Dr Emily Caseley (University of Leeds) for conducting molecular docking of drugs against the inhibitor-bound human KCNT1 structure.

## AUTHOR CONTRIBUTIONS

J.D.L. and L.M.D. conceived and designed the study and produced the draft manuscript. M.G.R., R.H., and Z.S. conducted and analyzed the *Drosophila* experiments. B.A.C. and G.Y.R. conducted and analyzed the electrophysiological experiments. J.D.L. designed and B.A.C. conducted molecular biology to generate the concatemeric constructs. K.J.S. conducted and analyzed the molecular docking. S.P.M. analyzed molecular structures and models. N.P. supervised and analyzed the research conducted by B.A.C.. All authors contributed to the manuscript preparation and reviewing. M.G.R., B.A.C, and R.H. contributed equally to this work and should be considered as co-first authors.

## DATA AVAILABILITY

The data that support the findings of this study are available from the corresponding authors, upon reasonable request.

