## Supplementary Information for "Identification of new KCNT1-epilepsy drugs by *in silico,* cell and *Drosophila* modelling"

### Supplementary Results

#### Supplementary Table 1: Parameters derived from Boltzmann analysis of WT, Y796H, and concatemeric $K_{Na}$1.1 currents.

Data are the mean ± SEM (of *n* number of cells provided in $5^{th}$ column) half-maximal activation voltages ($V_{½}$) and apparent gating charge (*z*, obtained from the slope $k = RT/zF$) the Boltzmann function fitted to conductance-voltage plots. Currents were recorded from CHO cells transiently transfected with plasmids to form homotetrameric WT or Y796H $K_{Na}$1.1 channels, or with concatemeric homomeric or heteromeric constructs. A hypthen “-“ between the subunits indicates the tethered tandem dimer (linker: GGGSGGGS) and the forward slash (/) between the subunits indicates that they are linked by the cleavable T2A motif (linker: GSGEGRGSLLTCGDVEENPG).

| KCNT1 construct | Linker | $V_{½}$ (mV) | *z* | *n* |
|---|---|---|---|---|
| WT | none | 7.38 ± 4.35 | 1.01 ±0.13 | 6 |
| WT-WT | Tethered dimer | -15.72 ± 1.85 | 1.46 ±0.17 | 5 |
| WT/WT | Cleavable T2A | -2.74 ± 7.69 | 0.89 ±0.07 | 5 |
| Y796H | none | -39.99 ± 2.27 | 0.92 ±0.08 | 6 |
| Y796H-Y796H | Tethered dimer | -50.83 ± 5.26 | 0.92 ±0.16 | 5 |
| Y796H/Y796H | Cleavable T2A | -42.19 ± 7.97 | 0.82 ±0.12 | 5 |
| WT/Y796H | Cleavable T2A | -16.42 ± 8.97 | 0.71 ±0.09 | 6 |
| Y796H/WT | Cleavable T2A | -21.51 ± 5.86 | 0.80 ±0.08 | 6 |

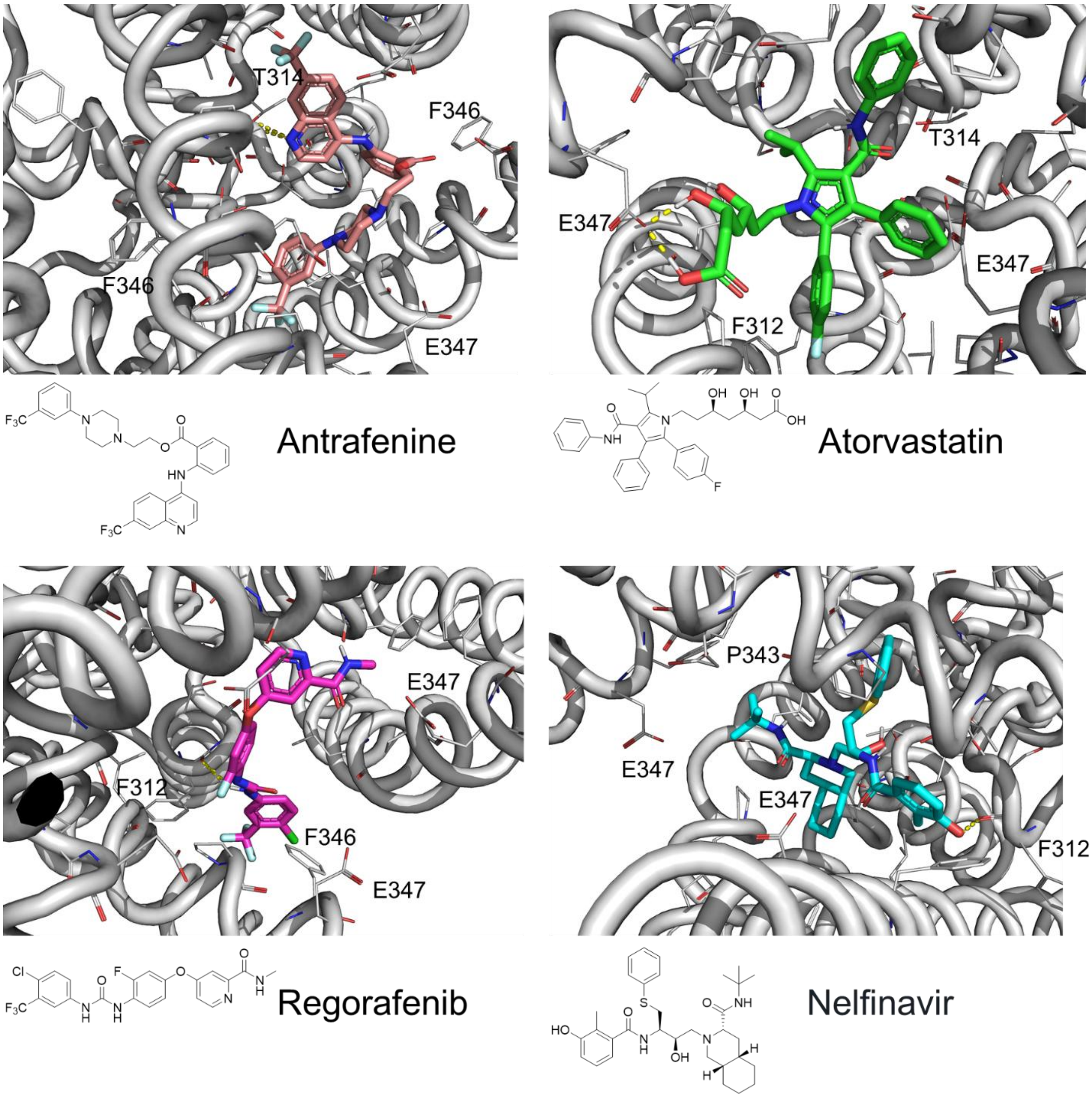


**Supplementary Figure 1: Predicted interactions of drugs with the $K_{Na}$1.1 pore domain (PDB: 5U70) though molecular docking in Glide (Schrödinger).**

**Supplementary Table 2. Glide (Schrödinger) docking scores for drugs interacting with the chicken activated KCNT1 channel structure (PDB: 5U70) and the inhibitor-bound human KCNT1 channel structure (PDB:8HKQ).**

| **Drug** | **Chicken** | **Human** |
|---|---|---|
| Antrafenine | -8.88 | -8.25 |
| Atorvastatin | -7.59 | -5.90 |
| Nelfinavir | -8.01 | -8.22 |
| Regorafenib | -7.88 | -8.32 |

**A**

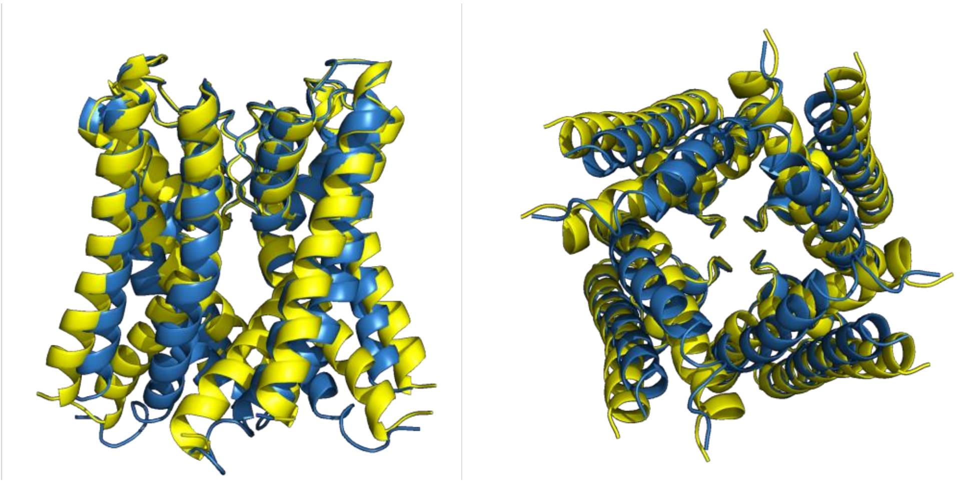

**B**

```
       <-------------S5------------->     <-----PH----->
Gga    ²⁴⁴SAMFNQVLILICTLLCLVFTGTCGIQHLERAGEKLSLFKSFYFCIVTFS
Hsa    ²⁶⁵SAMFNQVLILFCTLLCLVFTGTCGIQHLERAGENLSLLTSFYFCIVTFS

       <-SF->    <---------------S6------------->
Gga    TVGYGDVTPKIWPSQLLVVIMICVALVVLPLQFEELVYLWMERQKSGG³⁴⁰
Has    TVGYGDVTPKIWPSQLLVVIMICVALVVLPLQFEELVYLWMERQKSGG³⁶¹
```

**Supplementary Figure 2. Conservation in the pore domain between the chicken and human KCNT1 structures. A** Three-dimensional alignment of pore domains from the structures of chicken (yellow, PDB:5U70) and human (blue, PDB:8HIR) active KCNT1. The root mean square deviation between atoms is 1.3 Å. **B** Sequence alignment of the chicken (*Gga*) and human (*Hsa*) KCNT1 pore domains indicated in **A** comprising the S5 and S6 transmembrane segments, the pore helix (PH), and selectivity filter (SF) as indicated. Differences between species are highlighted in yellow.

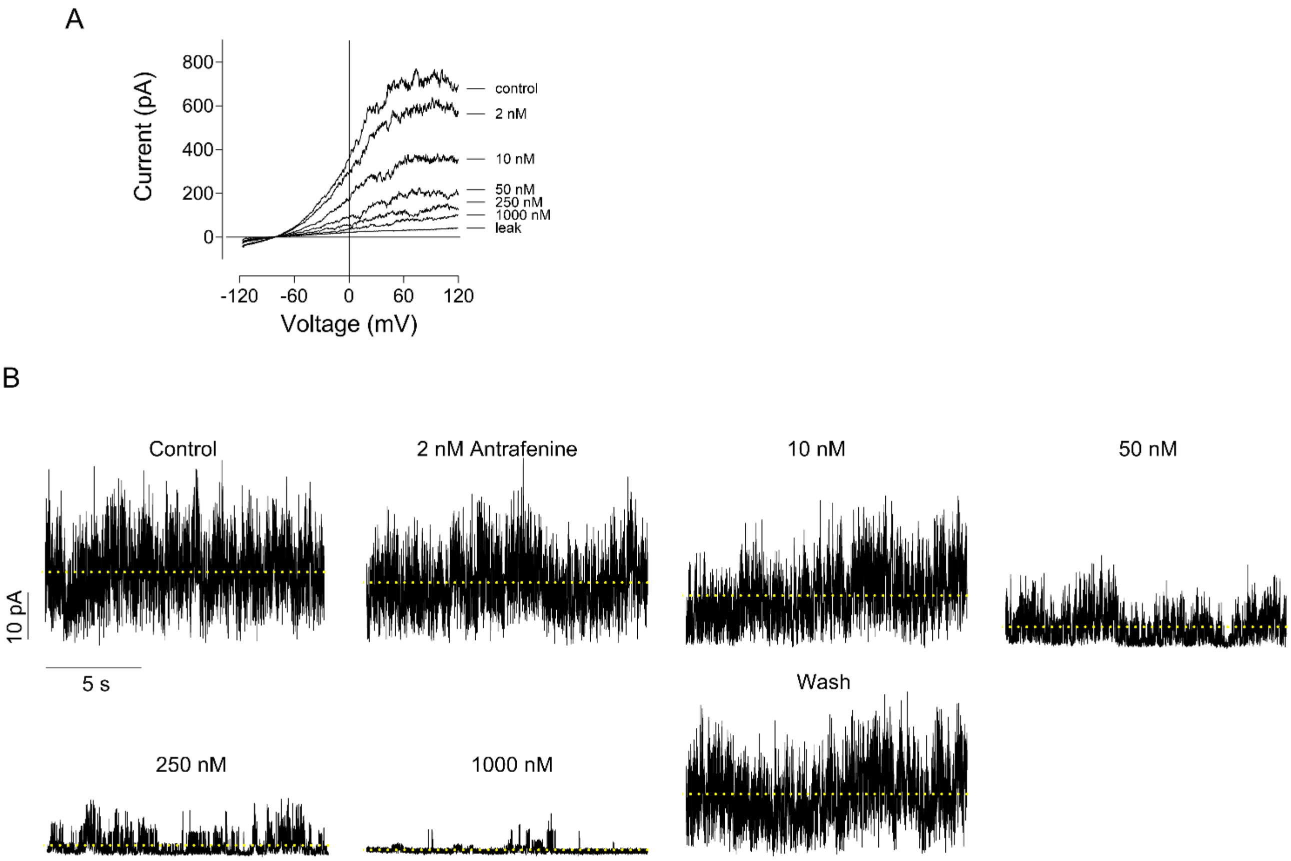


**Supplementary Figure 3. Examples of the current traces used to build concentration-response curves. A** Macroscopic G288S KCNT1 currents recorded in the inside-out patch in the presence of different concentrations of nelfinavir. Leak current was determined by replacing KCl in the bath solution with equimolar amount of NaCl. **B** Recording of a current in an inside-out patch containing 10 – 15 R398Q KCNT1 channels in the presence of different concentrations of Antrafenine, at 0 mV. The dashed lines represent the mean current amplitude of each trace.

**Supplementary Table 3**. One-way ANOVA results from the analysis of the Drosophila seizure phenotype data presented in Figure 6. ns - not significant.

| **G288S** | | | | | |
|---|---|---|---|---|---|
| | Antrafenine | Nelfinavir | Atorvastatin | Regorafenib | Bepridil |
| F | 44.78 | 52.39 | 1.529 | 0.7738 | 7.947 |
| P value | <0.0001 | <0.0001 | 0.2317 | 0.5799 | 0.0002 |
| P summary | **** | **** | ns | ns | *** |
| R squared | 0.8924 | 0.8881 | 0.2342 | 0.1621 | 0.6234 |

| **R398Q** | | | | | |
|---|---|---|---|---|---|
| | Antrafenine | Nelfinavir | Atorvastatin | Regorafenib | Bepridil |
| F | 25.10 | 68.90 | 3.880 | 8.160 | 4.379 |
| P value | <0.0001 | <0.0001 | 0.0192 | 0.0002 | 0.0069 |
| P summary | **** | **** | * | *** | ** |
| R squared | 0.8451 | 0.9323 | 0.4630 | 0.6602 | 0.5104 |

| **R928C** | | | | | |
|---|---|---|---|---|---|
| | Antrafenine | Nelfinavir | Atorvastatin | Regorafenib | Bepridil |
| F | 5.341 | 35.92 | 0.8976 | 1.970 | 8.246 |
| P value | 0.0001 | <0.0001 | 0.4932 | 0.1076 | <0.0001 |
| P summary | *** | **** | ns | ns | **** |
| R squared | 0.3022 | 0.7623 | 0.1108 | 0.2196 | 0.5076 |

**Supplementary Table 4.** P values obtained using One way ANOVA with Dunnett's multiple comparisons test of the data presented in Figure 6. Blue asterisks denote the statistically significant decrease in seizures compared to vehicle control, whereas red asterisks denote significant increase. ns – not significant. N is the number of independent experiments; number in brackets is the total number of flies analysed in each condition. For the controls, the number of independent experiments and the total number of flies for G288S, R398Q and R928C are 8 (88), 7 (141) and 12 (140), respectively.

| | G288S | | | | | R398Q | | | | | R928C | | | | |
|---|---|---|---|---|---|---|---|---|---|---|---|---|---|---|---|
| **[µM]** | **Antraf** | **Nelf** | **Atorv** | **Regor** | **Bepr** | **Antraf** | **Nelf** | **Atorv** | **Regor** | **Bepr** | **Antraf** | **Nelf** | **Atorv** | **Regor** | **Bepr** |
| **0.001** | <0.0001<br>****<br>N=4<br>(79) | 0.0052<br>**<br>N=4<br>(54) | 0.3267<br>ns<br>N=4<br>(136) | >0.9999<br>ns<br>N=2<br>(108) | 0.5514<br>ns<br>N=4<br>(85) | 0.0003<br>***<br>N=3<br>(62) | <0.0001<br>****<br>N=5<br>(261) | 0.0359<br>***<br>N=5<br>(113) | 0.0002<br>***<br>N=4<br>(135) | 0.5430<br>ns<br>N=4<br>(95) | 0.6765<br>ns<br>N=4<br>(54) | 0.9534<br>ns<br>N=12<br>(170) | 0.9998<br>ns<br>N=4<br>(59) | 0.9997<br>ns<br>N=5<br>(56) | 0.9923<br>ns<br>N=4<br>(46) |
| **0.01** | <0.0001<br>****<br>N=5<br>(143) | <0.0001<br>****<br>N=5<br>(71) | 0.1480<br>ns<br>N=4<br>(114) | 0.9929<br>ns<br>N=4<br>(54) | 0.9968<br>ns<br>N=4<br>(78) | <0.0001<br>****<br>N=6<br>(126) | <0.0001<br>****<br>N=4<br>(119) | 0.5236<br>ns<br>N=4<br>(122) | 0.0206<br>*<br>N=4<br>(57) | 0.3394<br>ns<br>N=4<br>(74) | 0.4387<br>ns<br>N=6<br>(102) | <0.0001<br>****<br>N=6<br>(115) | 0.9936<br>ns<br>N=4<br>(67) | 0.9865<br>ns<br>N=5<br>(56) | 0.0296<br>*<br>N=3<br>(57) |
| **0.1** | <0.0001<br>****<br>N=5<br>(101) | <0.0001<br>****<br>N=10<br>(153) | 0.7150<br>ns<br>N=2<br>(103) | 0.9009<br>ns<br>N=3<br>(145) | 0.0781<br>ns<br>N=3<br>(78) | <0.0001<br>****<br>N=5<br>(144) | <0.0001<br>****<br>N=2<br>(77) | 0.0676<br>ns<br>N=2<br>(89) | 0.0386<br>*<br>N=3<br>(175) | 0.0152<br>*<br>N=3<br>(66) | 0.0041<br>**<br>N=13<br>(216) | <0.0001<br>****<br>N=11<br>(177) | 0.9720<br>ns<br>N=4<br>(64) | >0.9999<br>ns<br>N=4<br>(55) | 0.0049<br>**<br>N=8<br>(110) |
| **1** | <0.0001<br>****<br>N=4<br>(56) | <0.0001<br>****<br>N=5<br>(73) | | 0.7301<br>ns<br>N=3<br>(64) | 0.0005<br>***<br>N=4<br>(65) | <0.0001<br>****<br>N=3<br>(71) | <0.0001<br>****<br>N=8<br>(113) | | 0.1793<br>ns<br>N=3<br>(70) | 0.0080<br>**<br>N=4<br>(55) | 0.0006<br>***<br>N=24<br>(301) | <0.0001<br>****<br>N=11<br>(161) | 0.3850<br>ns<br>N=8<br>(118) | 0.0360<br>*<br>N=5<br>(63) | 0.0002<br>***<br>N=9<br>(140) |
